# UNNT: A novel Utility for comparing Neural Net and Tree-based models

**DOI:** 10.1101/2023.09.12.557300

**Authors:** Venkata Vineeth Gutta, Satish Ranganathan Ganakammal, Sara Jones, Matthew Beyers, Sunita Chandrasekaran

## Abstract

Using deep learning in cancer research to tackle scientific challenges is becoming an increasingly popular technique. Advances in enhanced data generation, machine learning algorithms, and compute infrastructure have led to an acceleration in the use of deep learning in various domains of cancer research such as drug response problems. In our study, we explored tree-based models to improve the accuracy of a single drug response model and demonstrate that tree-based models such as XGBoost have advantages over deep learning models, such as a convolutional neural network (CNN), for single drug response problems. However, comparing models is not a trivial task. To make training and comparing CNNs and XGBoost (eXtreme Gradient Boosting) more accessible to users, we developed an open-source library called UNNT (A novel Utility for comparing Neural Net and Tree-based models). The case studies, in this manuscript, focus on cancer drug response datasets however the application can be used on datasets from other domains, such as chemistry.

**Author summary:** Advancement in data science, machine learning (ML), and artificial intelligence (AI) methods has enabled extraction of meaningful information from large and complex datasets that has assisted in better understanding, diagnosing, and treating cancer. The understanding of the drug response domain in cancer research has been accelerated with developing ML models to aid in predicting the effectiveness of the drugs based on a specific genomic molecular feature. In this study we developed a novel robust framework called UNNT (A novel Utility for comparing Neural Net and Tree-based models) that trains and compares deep learning method such as CNN and tree-based method such as XGBoost on the user input dataset. We applied this software to single drug response problem in cancer to identify the best performing ML method based on the National Cancer Institute 60 (NCI60) dataset. In addition, we studied the computational aspects of training each of these models where our results show that neither is evidently superior on both CPUs and GPUs while training. This shows that when both models have similar error rates for a dataset the hardware available determines the model choice for training as discussed in tables 7 and 11. UNNT: https://github.com/vgutta/UNNT.git

## Introduction

To leverage machine learning (ML) for cancer applications, the National Cancer Institute (NCI) at the National Institutes of Health (NIH) in collaboration with the Department of Energy (DOE) established the Joint Design of Advanced Computing Solutions for Cancer (JDACS4C) program [1]. This program developed three pilot projects focused on cancer research: Pilot 1-cellular level; Pilot 2-molecular level; Pilot 3-population level [2]. Along with the pilots, NCI-DOE also developed the CANDLE (Cancer Distributed Learning Environment) [3] project for hyperparameter optimization (HPO) on the models.

Our work in this paper is related to a subset of the drug response problems addressed in Pilot 1-cellular level. Specifically, our work builds on the existing single drug response predictor models officially known as P1B3, which uses a deep neural network to model tumor growth based on gene expression, drug concentrations, and drug descriptors data. We compared the performances of the existing CNN-based P1B3, built within the CANDLE framework, with the new tree-based methods. We show that a tree-based method, like XGBoost, is a better model than neural network CNN when the training data, such as drug response data, is tabular.

## Background

Unlike computer vision models, which rely on unstructured data such as images, the CANDLE framework drug response, and other models, rely on structured data in tabular format. The big breakthrough in Deep Learning came because of the ability of neural networks to perform well on unstructured data such as images. Deep neural networks have been successfully adapted to various domains outside of computer vision such as natural language processing (NLP) [4] and with the CANDLE framework to various problems in cancer research [3]. We tested the existing CNN model architecture used in CANDLE’s single drug response model with the NCI60 dataset and it peaked at an accuracy around 70%. Further improving such models will require data augmentation or changing to a new model architecture, and recent studies [5] have shown that tabular data may not require complex black box models such as CNNs to perform well. Gradient boosted tree models such as XGBoost [6] can match or exceed the performance of deep learning models [5]. State-of-the-art deep learning models for tabular data perform worse than XGBoost when tested on new data not in their respective original studies [5].

Tree-based models are supervised learning methods that create decision trees based on the training data provided. Decision trees are nodes in the tree-based model that create a “split” at a particular point in the data range for a particular feature in the data inferring rules during this process. They can be used for both classification and regression problems. During training, XGBoost builds decision trees sequentially and then uses a technique known as boosting where each successive tree gives more weight to examples that were previously misclassified. After each iteration, XGBoost computes the gradients of loss functions based on predictions and then creates a new decision tree to reduce errors made by previous trees [6].

The CNN model consists of a multilayer perceptron (MLP) and is a feed forward neural network. It generally consists of an input layer, hidden layers, and an output layer. The input layer receives the data and the hidden layers learn a continuous function based on the training data. These consist of convolutional layers called filters (kernels) that slide over the input data and compute the dot product between their weights and the input data. Following this operation, known as a convolution, non-linearity is introduced into the network using an activation function to learn complex relationships between features. Then a pooling layer, also using a kernel, slides across the data to reduce spatial dimensions and overfitting. A series of convolutional and pooling layers are typically followed by at least one fully connected layer to learn high-level features extracted by previous layers and the relationships between those features. After the forward pass through the network, a loss function is used to compare CNN’s output to the ground truth and is used to update the network’s weights and biases using backpropagation and gradient descent. Backpropagation computes the gradient of the loss functions for each of the weights and biases in the network going back to the input layer and these gradients are used to update the model parameters to minimize loss using optimization techniques such as gradient descent. Finally the output layers perform predictions which can be classification or a numerical output in a regression problem. For classification, a softmax layer converts the raw output from the network into class probabilities. Many other regularization techniques such as dropout are applied to prevent overfitting and improve model performance. This network architecture achieved state-of-the-art results in domains with grid-like unstructured data such as images but is not ideal for structured data.

## Design and Implementation

To provide users with structured (tabular) data the option to compare both the CNN and XGBoost models, we have developed an open-source comparative library called UNNT that allows users to bring their data to train both models. The XGBoost model relies on the open source libraries from Distributed (Deep) Machine Learning Community (DMLC) that UNNT uses to provide users the ability to train XGBoost models [6].

For data preprocessing, calculating metrics after training, and displaying results, we rely on other packages such as Pandas, Scikit-Learn, Numpy, Matplotlib.

To build CNN models we employ some of the functionality provided by the CANDLE library in Pilot 1 Benchmark 3 to do data preprocessing, model definition and instantiation, and model training. This has dependencies such as TensorFlow 1.0. We also use scikit-learn for metrics to test the model performance.

The preprocessing steps are important for the predictive accuracy of our pretrained models to work on any data users bring to train CNN and XGBoost models. Because the exact format of user provided datasets is unknown, users are responsible for any preprocessing and data cleaning steps that would be necessary prior to model training. Users need to decide what features they want to keep in the dataset being used to train and test the models and our models should be able to accommodate data of any shape as long as it fits into the device memory.

Once users provide their data, UNNT splits that data into training, validation, and testing sets and the user can specify the percentage of data used for testing and validation. We convert each of the data splits into numpy arrays to train using both CNN and XGBoost. There are several hyperparameters that can be set, for both CNN and XGBoost models, and we will have recommended default parameters which can be customized by the user. Users have full control of the parameters tested to find the best combination of hyperparameters for their dataset for each of the models.

Finally UNNT provides users error metrics to evaluate both models trained using their data including R^2^ and Root mean square error (RMSE).

## Data

The data for the study was obtained from The Predictive Oncology Model and Data Clearinghouse (MoDaC) [7] data warehouse that was released as part of JDACS4C [1]. The dataset includes RNA-Seq expression profiles, drug response and molecular drug descriptors for National Cancer Institute 60 (NCI60) [8] cell lines.

The RNA-Seq expression data from these various cell lines are normalized using ComBat-seq [9]. Drug dose response data obtained from these cell lines are normalized using a Hill slope model with three bounded parameters [10] and drug molecular descriptors were generated using Dragon software package (version 7.0) [11].

To compare model performances of XGBoost and CNN models, we extracted train, validation and test datasets corresponding to the NCI60 cell lines from our combined datasets that include gene expression values [12], drug response values [13], and drug descriptors [14] found in MoDaC [7].

Our NCI60 data consists of data similar to the NCI60 data the original Single Drug Response model in CANDLE Benchmarks was trained with, with two main differences. Firstly our dataset uses only using lincs1000 genes [15] in the RNA sequence gene expression data instead of all the protein coding genes, due to the importance of those genes in the dataset. Secondly, the original NCI60 dataset in drug response data uses ‘Growth’ and ‘Concentration’ values where ‘Growth’ is the target variable and ‘Concentration’ is a feature which we no longer use. ‘Growth’ is replaced with ‘AUC’ data in the drug response dataset because area under the curve (AUC) [16] is a more definitive parameter that combines both potency (concentration) and efficacy of a drug and is more robust when comparing a single drug across multiple cell lines for similar dose levels.

## Installation + running instructions

1. Download or git clone UNNT repository from GitHub: https://github.com/vgutta/UNNT.git
  - *git clone https://github.com/vgutta/UNNT.git*
  - *cd UNNT*
2. Prerequisites
  a. Anaconda package manager: Installation instructions for your specific system can be found here
3. Install and activate the relevant Anaconda environment
  a. NVIDIA GPU
    - *conda env create -f gpu environment.yml -n UNNT gpu*
    - *conda env activate UNNT gpu*
  b. CPU
    - *conda env create -f environment.yml -n UNNT*
    - *conda activate UNNT*
4. Default usage trains CNN and XGBoost models on NCI60 datasets
  a. *cd UNNT*
  b. CPU: *python3 unnt.py*
  c. GPU: *python3 unnt.py --gpu*
5. Data
  a. All data used to train models with UNNT must be placed in the **data** folder at the root of this repository. The data necessary to run the default settings using NCI60 datasets are also located in the **data** folder. Provide the name of the file for the configuration parameter **data file**. Name of file must be in quotes and is case-sensitive. The dataset provided is required to be in csv format.
  b. Provide data that is cleaned and preprocessed into the final format necessary for training. The columns in the dataset must be features. Provide the **target variable** the models train on by setting the **target variable** configuration parameter in **/UNNT/tree config**.**txt**
6. Configuration files
  a. In addition to configuration on data necessary for the entire software, **/UNNT/tree config**.**txt** also contains XGBoost model parameters.
  b. **/UNNT/cnn config**.**txt** contains the configurations specific for CNN model. Many of these are specific to NCI60 dataset used for demo in this software and thus won’t affect the models trained using custom data.

## Results

### Experimental Setup

The compute resources for this work used NSF sponsored cluster, DARWIN, [17] at UDEL and Perlmutter [18], at LBNL. DARWIN and Perlmutter have GPU and CPU nodes. DARWIN uses NVIDIA V100 GPUs while Perlmutter has the latest A100 GPUs. DARWIN has AMD EPYC 32 core processors which are similar to Perlmutter that has an AMD EPYC 7713 64-core CPU.

### XGBoost

We trained an XGBoost model using NCI60 expression, dose response, and drug descriptor data with AUC as the target (predictor) variable. Our observed test accuracy yielded 0.83 for R^2^ error and 0.05 RMSE error. In addition, we created a separate dataset set aside before training and included 10% of the cell lines. Table 1 shows the difference in test errors between the two.

**Table 1.**
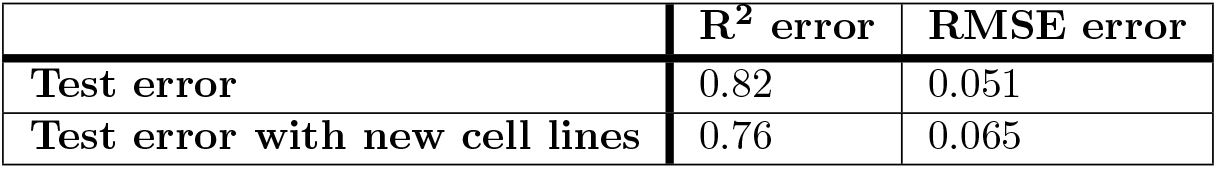
XGBoost Errors for model trained NCI60 data.

To find the best set of hyperparameters, we performed hyperparameter optimization using grid search technique and cross validation. Grid search trains a new model for every combination of hyperparameters while cross validation uses a different subset as test data to get an average across five subsets. Best set of hyperparameters found were ETA:0.1, Max depth: 10, Subsample: 0.5, N estimators:500. We used these hyperparameters to train a new model and the results are shown in Table 1.

Hyperparameter optimization led to slight improvement that was lower than our initial expectations. This was a result of well documented ranges for the various parameters and thus we happened to choose values that were close to optimal for each of the hyperparameters.

XGBoost model training requires the training data to be fully merged before training commences and this resulted in memory issues. To solve the memory issues, we experimented with smaller datasets and reduced the number of drugs from 30,000 to 159, based on an FDA approved drugs list [13].

### CNN

We trained the original CNN model using the new NCI60 data as described in the data section above. The results shown in Table 3 were after applying hyperparameter optimization (HPO) and to determine the best parameters where the learning rate is 0.01 and tanh as the activation function. The model failed to converge on other activation functions such as ReLU for NCI60 data.

**Table 2.** XGBoost Errors for model trained on NCI60 data with only FDA approved drugs list.

|  | R<sup>2</sup> error | RMSE error |
| --- | --- | --- |
| Test error | 0.84 | 0.069 |
| Test error with new cell lines | 0.66 | 0.094 |

**Table 3.** CNN errors for NCI60 after training after performing HPO.

|  | R<sup>2</sup> error | RMSE error |
| --- | --- | --- |
| Test error | -30.32 | 0.81 |

As shown in in Table 3, the CNN model performed much worse than XGBoost when trained on NCI60 data. The RMSE metric’s best possible value is 0 and it can go to infinity, but a value such as 0.81 does not give us good insight into the quality of the regression model. On the other hand, the negative R^2^ value for the test error indicates the model performance is poor but its magnitude does not indicate how poorly it performed. The best value for R^2^ is 1, meaning the model completely explains predicted data variability [19]. These results indicate that both R^2^ and RMSE metrics show XGBoost model is outperforming the CNN model trained on this NCI60 dataset.

### Training times of CNN and XGBoost

The original CNN model is capable of running on both GPUs and CPUs, by design, since it is built with the TensorFlow framework. While the XGBoost model runs on CPU by default it can also be trained on GPU where we use the parameter *tree method=“gpu_hist”* in the XGBRegressor function. This means both models in UNNT can be accelerated using GPUs. In the following section we show comparisons of model training times with CPUs and GPUs.

#### Training of full NCI60 drugs

In addition to the model built using only the FDA approved drugs, we also built an XGBoost model using all the available drug data we have access to, however, use of 30,000 drugs presented many challenges due to the volume. The main challenge using the entire drug list entailed finding a system with at least 500GB of memory for the merged data before we train the XGBoost model.

CNN models can have varying training times based on parameters used for training such as subsampling with fewer features, specifying fewer training, validation, test steps, and by reducing the number of training epochs. We discuss some of those results. Table 5 shows that the CNN model does not improve as the number of epochs increases, eliminating a benefit of higher epochs. And the difference between training times of CNN and XGBoost is large. Table 6 shows that CNN model converges to its optimal learning capacity in 1 epoch, hence it would still take three times longer to train than an XGBoost model trained with a V100 GPU in the best case scenario of 1 epoch.

**Table 4.** Results using XGBoost. Times for threads represents model runs on CPU with the corresponding threads used for speedup. Last row corresponds to running the same model on single NVIDIA V100 GPU.

| Threads | Time (hours) |
| --- | --- |
| 1 | 5.6 |
| 2 | 6 |
| 4 | 5.94 |
| 8 | 6 |
| 16 | 5.9 |
| 32 | 6.7 |
| 64 | 7.2 |
| V100 GPU | 0.3 |

**Table 5.** CNN model trained with all features on an NVIDIA V100 GPU.

| Epochs | R<sup>2</sup> error | RMSE error | Time (hours) |
| --- | --- | --- | --- |
| 1 | -30.46 | 0.821 | 2.3 |
| 5 | -30.32 | 0.819 | 11.4 |
| 10 | -30.32 | 0.819 | 22.96 |
| 15 | -30.32 | 0.819 | 34.36 |

**Table 6.** CNN model trained with all features on a CPU.

| Epochs | R<sup>2</sup> error | RMSE error | Time (hours) |
| --- | --- | --- | --- |
| 1 | -30.35 | 0.81 | 1.1 |
| 5 | -30.31 | 0.81 | 5 |
| 10 | -30.32 | 0.81 | 9.97 |
| 15 | -30.32 | 0.81 | 15 |

When comparing training times from Tables 5 and 6 we can see that the CNN model takes half as long to train on CPU compared with training on GPU. This is most likely a result of the size of the dataset where data transfer from CPU to GPU becomes a bottleneck and increases training time.

#### Training on FDA approved drugs

Table 7 shows us that an XGBoost model trains much faster on a GPU when training on a dataset that only contains FDA drug subset. We observed that the training times on the CPU increased as more threads were added. This occurs when the communication overhead is greater than the computational benefit of distributing a model across cores.

**Table 7.**
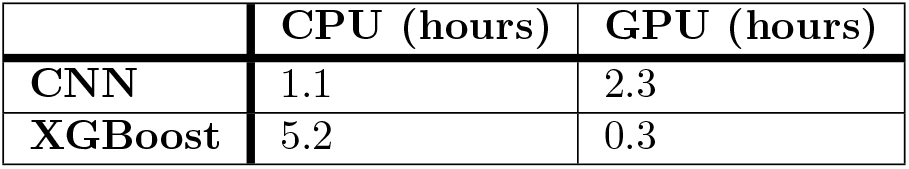
Fastest training times for CNN and XGBoost on CPU and GPU (all features)

Table 8 shows the one instance where CNN model trains faster than an XGBoost model when training on a similar dataset. Comparing Tables 8 and 9 we see the CNN model trains faster on a CPU even with less training data. The CPU results for CNN are not broken down by the number of threads because TensorFlow 1.0, the framework used to build CNN model, does not support threading on CPUs.

**Table 8.** Results using XGBoost. Times for threads represents model runs on CPU with the corresponding threads used for speedup. Last row corresponds to running the same model on single NVIDIA V100 GPU.

| Threads | Time (seconds) |
| --- | --- |
| 1 | 1495.44 |
| 2 | 1527.33 |
| 4 | 1534.78 |
| 8 | 1549.24 |
| 16 | 1568.54 |
| 32 | 1599.01 |
| 64 | 1639.83 |
| V100 GPU | 160 |

**Table 9.**
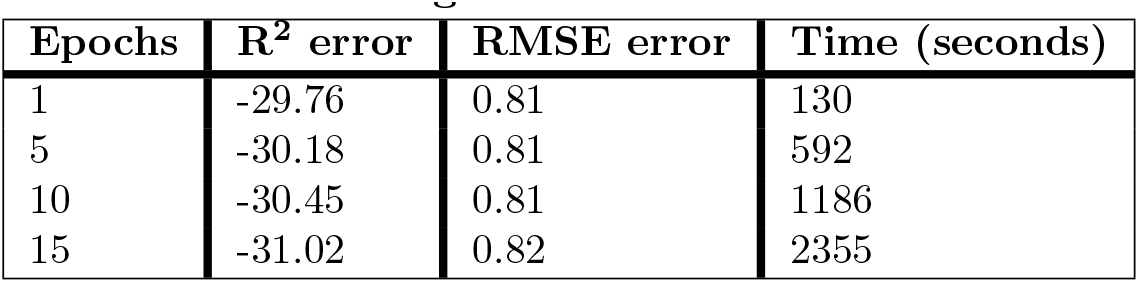
CNN model FDA drugs trained on a CPU.

| Epochs | R<sup>2</sup> error | RMSE error | Time (seconds) |
| --- | --- | --- | --- |
| 1 | -29.76 | 0.81 | 130 |
| 5 | -30.18 | 0.81 | 592 |
| 10 | -30.45 | 0.81 | 1186 |
| 15 | -31.02 | 0.82 | 2355 |

Tables 7 & 10 show the advantage of training XGBoost on a V100 GPU that is consistently the fastest for the same data.

**Table 10.** CNN model FDA drugs trained on a V100 GPU.

| Epochs | R<sup>2</sup> error | RMSE error | Time (seconds) |
| --- | --- | --- | --- |
| 1 | -30.07 | 0.81 | 300 |
| 5 | -30.96 | 0.81 | 1356 |
| 10 | -30.33 | 0.81 | 2672 |
| 15 | -31.35 | 0.81 | 5348 |

**Table 11.** Fastest training times for CNN and XGBoost on CPU and GPU (FDA model)

|  | CPU (seconds) | GPU (seconds) |
| --- | --- | --- |
| CNN | 592s | 1356s |
| XGBoost | 1495s | 160s |

## Conclusion

Exploring a niche domain such as drug response modeling for cancer cell lines, we show that utilizing a neural network (CNN) does not yield the best results. Instead, we show the impact of tree-based XGBoost model over a CNN model especially when the datasets trained on are tabular and running on a GPU. Our results demonstrate that using the same dataset, an XBoost model is faster than a CNN model while running on an NVIDIA GPU. An observable downside to using XGBoost is the larger memory requirement for training, as documented in this work, and this varies depending on the size of the dataset. We have also developed a utility, UNNT, that allows users to bring their data and build models such as CNNs and XGBoost as well as compare how the models perform on the given dataset. Thus UNNT would be a useful utility for domain scientists to experiment with two unique model architectures for tabular data.

## Future Work

As part of future work, we will incorporate other machine learning model architectures such as autoencoders to augment CNN and XGBoost by encoding certain classes of features, such as gene expression values. Increasing the number of available evaluation metrics will be useful for a wider range of domains and the addition of classification models will necessitate a wider range of evaluation metrics. In addition, there are many features such as hyperparameter optimization that augment UNNT’s existing capabilities even though users can currently adjust and refine hyperparameters of their models manually. Adding functionality to reduce the preprocessing and data cleaning done by the user will allow UNNT’s capability to broaden even further. The prevalence and growth of tabular data going forward will only elevate the need for software like UNNT.

## Acknowledgement

This work has been funded in whole or in part with Federal funding by the NCI-DOE Collaboration established by the U.S. Department of Energy (DOE) and the National Cancer Institute (NCI) of the National Institutes of Health, Cancer Moonshot Task Order No. 75N91019F00134 and under Frederick National Laboratory for Cancer Research Contract 75N91019D00024. This work was performed under the auspices of the U.S. Department of Energy by Argonne National Laboratory under Contract DE-AC02-06-CH11357.

---

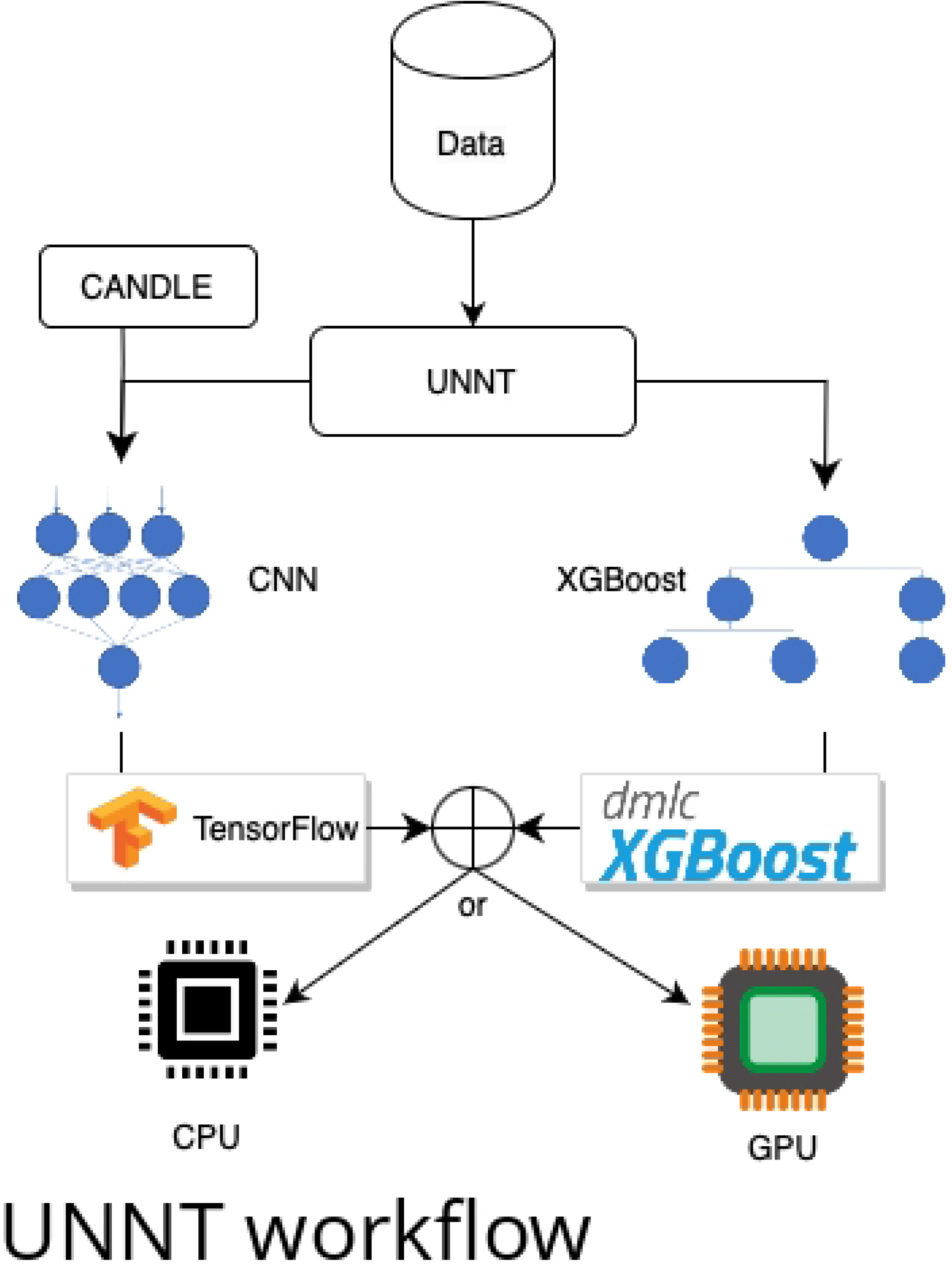

## Notes

### Competing Interest Statement

The authors have declared no competing interest.

